# Population Pharmacokinetics and Probability of Target Attainment Analysis of Vancomycin Following Intermittent and Continuous Infusion in Adults with Cystic Fibrosis

**DOI:** 10.1101/2025.06.26.661675

**Authors:** Manav Jain, Rachel Hudson, Zubin Bhakta, David C. Young, Venkata K Yellepeddi

**Author notes:** Manav Jain and Rachel Hudson contributed equally to this work. Author order was determined alphabetically.

## Abstract

Vancomycin is the drug of choice for treating pulmonary infections caused by methicillin-resistant *Staphylococcus aureus* (MRSA) in people with cystic fibrosis (PwCF). This study characterized the pharmacokinetics (PK) of continuous and intermittent vancomycin infusions following a loading dose using population PK (PopPK) modeling to inform dosing in PwCF. The PopPK model was developed using therapeutic drug monitoring (TDM) data from adult PwCF who received a vancomycin loading dose followed by intermittent, continuous, or both infusion types for MRSA-related pulmonary exacerbations. A total of 212 samples were collected following 90 intermittent and 42 continuous infusions in 21 patients. The final model was a two-compartment model with first-order elimination, incorporating creatinine clearance (CrCL) as a covariate on vancomycin clearance (CL). The estimated CL and volume of distribution were 4.05 L/h/70 kg and 22.5 L/70 kg, respectively. The model was used to predict the probability of target attainment (PTA) following a single intermittent loading dose (500–1500 mg) and continuous infusion (500–6000 mg) over 24 hours. PTA was assessed using efficacy and toxicity thresholds defined by Area Under the Curve_0-24_ (AUC_0-24_)/Minimum Inhibitory Concentration (MIC) ratios ≥400 mg·h/L and <650 mg·h/L, respectively. At a MIC of 1 µg/mL, a loading dose of 500 mg followed by a 3750 mg continuous infusion achieved PTA targets for efficacy (66.7%) and safety (82.7%). These findings support the use of PopPK modeling to guide vancomycin dosing strategies for MRSA pulmonary infections in PwCF.

## Introduction

A vast majority of pulmonary exacerbations in PwCF are caused by MRSA infections (1, 2). Unfortunately, MRSA infections in this patient population are associated with a 27% higher mortality risk and a 6.2-year shorter life span compared to those without MRSA (3). In the United States, the prevalence rate of MRSA among PwCF increased from 2% in 2001 to 26% in 2013, with a more recent prevalence of 13% in 2024 (4). The American Thoracic Society recommends intravenous (IV) vancomycin as first-line therapy for treating MRSA infections in PwCF (1). However, despite widespread use, IV vancomycin dosing in PwCF is largely empirical and inconsistent (5). A national practice survey conducted across various CF Foundation (CFF) accredited care centers and affiliate programs in the U.S. revealed significant variability in vancomycin dosing, with reported regimens in adults ranging from 500 mg to 2000 mg every 6 to 12 hours. The absence of established dosing guidelines contributes to the wide variability in vancomycin regimens for this patient population. The survey highlighted the urgent need for optimized IV vancomycin dosing and TDM strategies to support more individualized, evidence-based care (6).

Vancomycin is a tricyclic glycopeptide antibacterial drug used to treat serious gram-positive bacterial infections, especially for MRSA (7). It has a narrow therapeutic index and exhibits large interindividual PK variations. Vancomycin exhibits 10-50% plasma protein binding, a half-life (t_1/2_) of 4-6 hours in patients with normal renal function, and is primarily eliminated unchanged in urine more than 80% recovered within 24 hours of IV administration (8, 9). Vancomycin has a CL of 0.058 L/kg/h and V_d_ between 0.4 to 1 L/kg (9). Accurate dosing is important to understand the efficacy and toxicity levels of vancomycin as nephrotoxicity and ototoxicity are the major dose-limiting factors.

Traditionally, vancomycin was titrated based on serum trough levels of 15 to 20 mg/L as an indicator of efficacy to treat chronic MRSA infections in PwCF (5). However, vancomycin trough concentrations of ≥15 mg/L significantly increase the risk of nephrotoxicity. An AUC/MIC ratio between ≥400-650 mg*h/L is considered optimal for efficacy, with higher exposures potentially increasing toxicity risk (10, 11). The AUC/MIC ratio is well established in individuals without CF, but its utility in PwCF remains uncertain due to significant PK alterations in this population.

Pathophysiological changes in PwCF contribute to alterations in the PK profiles of certain antibiotics through a combination of physiological and disease-related factors. Some of these factors include decreased plasma protein binding, increased renal clearance, increased glomerular filtration rate, chronic inflammation, and differences in tissue penetration (12). Therefore, characterizing the PK for antibiotic agents, including vancomycin among PwCF, is critical to recommend optimal dosing for maximal therapeutic efficacy and minimal toxicity. Data is scarce regarding the PK of vancomycin in adult PwCF. Pleasants et al. reported PK parameters of vancomycin in PwCF after a single IV dose of 15 mg/kg in 10 adult PwCF (13). More recently, the PK of vancomycin in PwCF after intermittent or continuous dosing were reported (14, 15). We recently reported the PopPK of vancomycin after intermittent dosing in adults PwCF (16). However, in a clinical setting where both intermittent and continuous infusion are prescribed, the characterization of vancomycin PK is necessary.

Both intermittent and continuous infusion of vancomycin have been prescribed in clinical settings, with no established evidence of preference for one dosing strategy over the other (17). While intermittent vancomycin infusion has traditionally been a standard approach, continuous infusion of vancomycin is gaining interest as an alternative dosing strategy, especially when AUC/MIC target attainment and reduced nephrotoxicity are desired. Chu et al., 2020, found that the vancomycin target concentration was easier to achieve with less risk of nephrotoxicity with continuous infusion (18). This approach may be particularly useful in patients at high risk of nephrotoxicity, such as those on other nephrotoxic drugs (e.g. aminoglycosides or loop diuretics), with septic shock, or requiring high-dose vancomycin. Continuous vancomycin infusion may increase the chances of achieving pharmacodynamic (PD) targets of AUC_0-24_/MIC ≥400 mg*h/L (19). Despite its theoretical advantages, continuous vancomycin infusion remains understudied in adult PwCF, highlighting the need for further clinical research.

To address the current gap in dosing guidance when vancomycin is given as intermittent or continuous infusion or both, the present study aims to characterize the PK of vancomycin administered via intermittent and continuous infusion in adult PwCF. A PopPK model was developed based on TDM and clinical data collected at the University of Utah Hospital.

Additionally, PTA was evaluated across different dosing scenarios using Monte Carlo simulation to assess both efficacy (AUC_0-24_/MIC ≥400 mg*h/L) and safety (AUC_0-24_/MIC <650 mg*h/L) targets. These findings can help guide vancomycin dosing following a loading dose to optimize treatment of MRSA pulmonary infections in adult PwCF.

## Results

### Patient characteristics

212 samples were collected after administering 90 intermittent and 42 continuous doses of vancomycin to 21 patients. All vancomycin serum concentrations were quantified above the LLOQ of 1.1 mg/L. Demographic and clinical data collected in this study are shown in **Table 1**. 61.1 % of patients received a 750 mg intermittent infusion while 21.4 % of patients received 2500 mg of continuous infusion (**Supplementary Table 1A and 1B**).

**TABLE 1.** Demographic and clinical characteristics of patients.

| Variable | Mean ± SD | Median (range) | N (%) |
| --- | --- | --- | --- |
| Gender |  |  |  |
| Male | N/A | N/A | 5 (23.8) |
| Female |  |  | 16 (76.2) |
| Concomitant inhaled antibiotics |  |  |  |
| Inhaled tobramycin | N/A | N/A | 12 (57.1) |
| Inhaled aztreonam |  |  | 3 (28.6) |
| Both |  |  | 6 (14.3) |
| Race |  |  |  |
| White | N/A | N/A | 20 (95.2) |
| Hispanic or Latino |  |  | 1 (4.8) |
| Age (years) | 29.3 ± 11.4 | 27 (19-60) | N/A |
| Weight (kg) | 61.8 ± 16.6 | 54.5 (43.6-105.7) | N/A |
| Baseline |  |  |  |
| Serum creatinine (mg/dL) | 0.76 ± 0.22 | 0.7 (0.44-1.45) | N/A |
| Creatinine clearance (mL/min) | 124.9 ± 34.2 | 122.7 (42.8-214.8) | |
| BSA normalized creatinine clearance (mL/min/1.73 m<sup>2</sup>) | 72.2 ± 19.8 | 71 (24.8-124.2) | |
| Day 7 |  |  |  |
| Serum creatinine (mg/dL) | 0.69 ± 0.17 | 0.65 (0.43-1.17) | N/A |
| Creatinine clearance (mL/min) | 137.8 ± 33.5 | 137.6 (51.2-198.9) | |
| BSA normalized creatinine clearance (mL/min/1.73 m<sup>2</sup>) | 78.5 ± 19.3 | 79.6 (29.6-115) | |
| Vancomycin infusion |  |  |  |
| Intermittent | N/A | N/A | 7 (33.3) |
| Continuous |  |  | 7 (33.3) |
| Both |  |  | 7 (33.3) |
| Vancomycin intermittent dose (mg) | 855.6 ± 157 | 750 (500 – 1500) | 90 (68.2) |
| Vancomycin continuous dose (mg) over 24 hours | 3161 ± 1798 | 2500 (1500-9000) | 42 (31.8) |
| Day of starting continuous dose after intermittent dose* | 5.9 ± 3.1 | 6 (2 -14) | N/A |
| Sample collection after continuous infusion (h) | 17.9 ± 6.1 | 16.8 (7.2 – 33.1) | N/A |

h=hour; kg=kilograms; mg/dL=milligram per deciliter; min=minute; mL=milliliter; mL/min=milliliter per minute, \*Only in patients who received both intermittent and continuous vancomycin

### Population pharmacokinetic model

Both one- and two-compartment models with first-order elimination were evaluated using different residual error structures, including additive, proportional, and combined models. The model also incorporated the rate and duration of the intravenous infusion for each patient. A two-compartment model with first-order elimination and a proportional residual error structure was selected based on model diagnostics and PK parameters.

The population parameter estimates from the final model are shown in **Table 2**. Variability was lowest for the central volume of distribution (V1) and highest for the peripheral volume of distribution (V2), indicating more uncertainty in the estimation of the peripheral volume of distribution. IIV was included in CL. Although both weight and CrCL significantly affected CL (*P* < 0.05) and the reduction in OFV and IIV was higher for CrCL than for weight, multivariate analyses indicated including both weight and CrCL covariates didn’t improve the model over individual covariates. Hence, in the final model, CrCL was retained as a predictor of clearance as it produced the most significant minimization of the OFV (Δ28.71) and reduced the IIV. A summary of the effects of sequential covariates is provided in **Supplementary Table 2**.

**TABLE 2.** Final NONMEM Parameter Estimates and Bootstrap Results.

|  |  |  | NONMEM | Bootstrap (n = 1,000) |  |  |
| --- | --- | --- | --- | --- | --- | --- |
| Structural model parameters |  |  | Estimate | Median | 95% CL | CV (%) |
| CL<br>(L/h) | $\exp(\Theta_1)$ | Clearance | 4.05 | 3.91 | (1.73 – 4.33) | 16.70% |
| V1 (L) | $\exp(\Theta_2)$ | Central volume | 22.5 | 22.5 | (18.9 – 25.9) | 6.40% |
| Q (L/h) | $\exp(\Theta_3)$ | Intercompartmental<br>clearance | 1.98 | 2.1 | (0.65 – 3.84) | 31.30% |
| V2 (L) | $\exp(\Theta_4)$ | Peripheral volume | 67.6 | 72 | (45.0 – 1177.9) | 139.50% |
| Covariate effect parameters |  |  |  |  |  |  |
| CL ~<br>CRCL | $\Theta_6$ | Creatinine clearance<br>effect on clearance | 0.637 | 0.703 | (0.406 – 2.294) | 46% |

95% CL, confidence level reported as 2.5 and 97.5% confidence limits; CV, coefficient of variation calculated from mean bootstrap parameter estimates. CL=clearance; CL CRCL=creatinine clearance; L=liter; L/h=liter/hour

Diagnostic plots were generated to assess model fit between observed vancomycin concentrations and population-predicted and individual-predicted values. The conditional-weighted residuals versus the population-predicted vancomycin concentration plots were also evaluated (**Supplementary Figure 1**). Residual plots showed that the residual remained mainly centered around zero with no systematic or time trends throughout the dosing interval (CWRES vs PRED and CWRES vs TIME). The observed plasma concentrations versus population-predicted and individual-predicted plasma concentrations (DV vs PRED and DV vs IPRED) were evenly distributed around the line of unity, suggesting that there was no major bias in the structural model. Overall, visual inspection revealed that the final model described the data more tightly than the base model.

In the prediction-corrected visual predictive check (pcVPC) plot, the median observed concentrations closely followed the model-predicted median and ranged fairly within 95% PI around the predicted median, suggesting good central tendency. Additionally, the 5^th^ and 95^th^ percentiles of the observed data fell within 95% PI of the simulated corresponding percentiles, suggesting the model adequately captured the variability in the observed data. The model also captured the changes in vancomycin exposure over time, both in observed and predicted concentrations (after 200 hours) (**Figure 1)**. Numerical assessment diagnostics showed that at 95% PI, the proportion of observations outside the simulated PI was low, with 0.94% (False Positive (FP)) of observed values above and 0.47% (False Negative (FN)) of values below the upper and lower bounds, respectively. As PI decreased, the percentage of FP and FN (at 90% PI, FP = 1.89%, FN = 2.36%) increased, suggesting a little underprediction at narrower PI. Overall, the coverage was within the acceptable range. Thus, the pcVPC validates the appropriateness of the model in capturing the typical trend as well as variability in the observed PK data.

**Figure 1.**
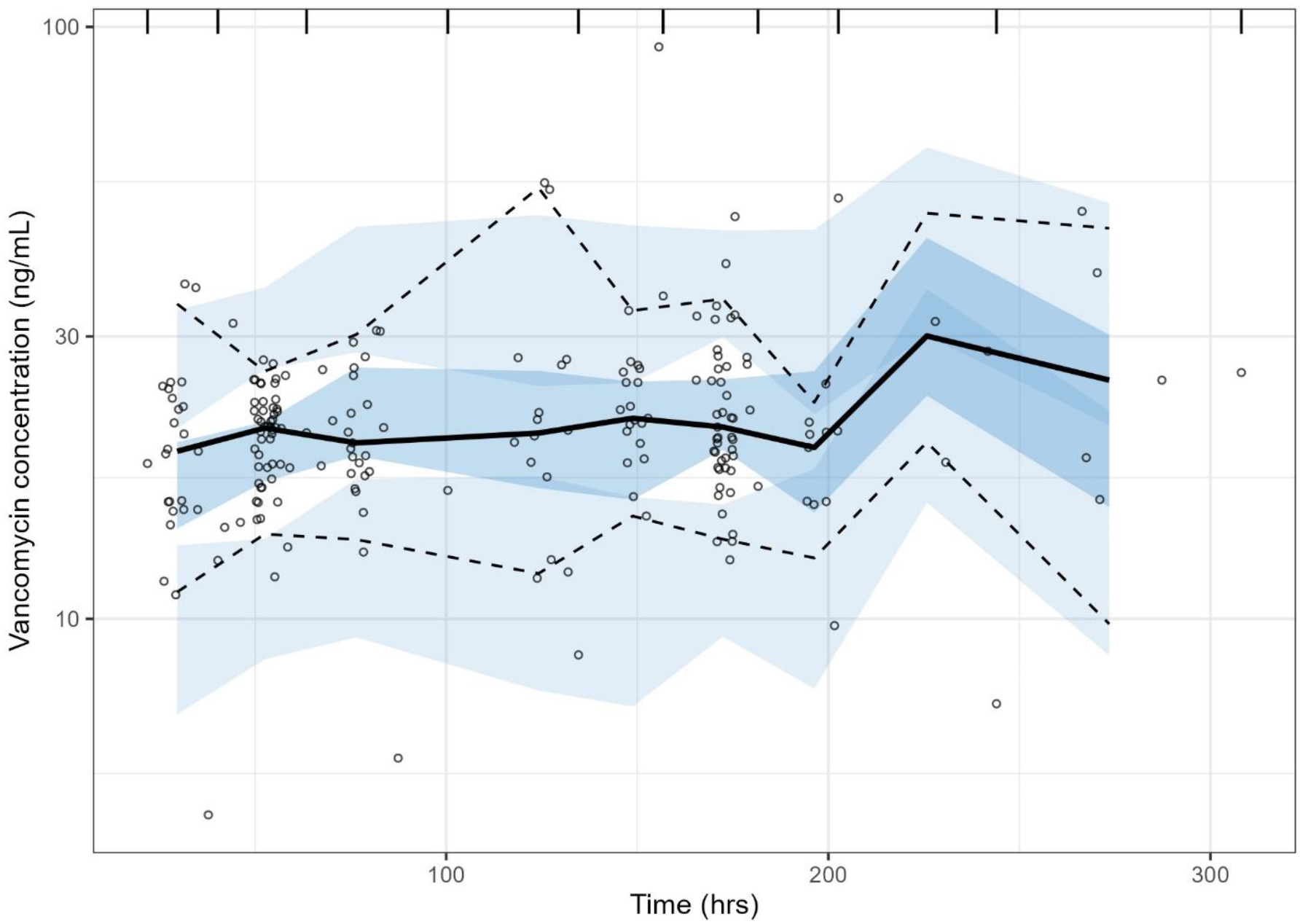
Prediction-corrected visual predictive check for the final model. Black dots observed values; solid black line, medial (50^th^ percentile); dashed black lines, 5^th^ and 95^th^ percentiles for the observed values; dark and light blue shaded regions, 95% confidence intervals for the 5^th^, 50^th^, and 95^th^ percentiles of the simulated data. ng/mL=nanogram/milliliter

The PTA in plasma across different vancomycin dosing scenarios (loading dose intermittent infusion followed by continuous infusion) was evaluated with the target AUC_0-24_/MIC ≥400 & < 650 mg*h/L and AUC_0-24_/MIC ≥650 mg*h/L. The AUCs after the first 2 days of initial dosing (AUC_0–24_ and AUC_24–48_) following single loading dose of 500, 750, 1000, 1250, and 1500 mg, along with various continuous dosing infusions, were used for PTA analysis. **Figure 2 (A & B)** shows the percentage of patients achieving the efficacy target AUC_0-24_/MIC (MIC = 1 µg/mL) ratios across all dosing scenarios simulated from the PopPK model. The PTA was highest with continuous infusion of 3750 mg (66.7 %) and 2750 mg (57.3 %), and the loading dose was 500 mg for AUC_0-24_ and AUC_24-48_, respectively. At the highest efficacy target, the PTA for safety was achieved in 82.7 % and 77.4 %. When the toxicity targets were considered, the PTA increased with increasing continuous dosing infusion across all the loading dose scenarios **(Supplementary Table 3 A-D)**. The box plots show AUC_0-24_ and AUC_24-48_ versus various continuous infusions along with a single loading dose of 500, 750, 1000, 1250 and 1500 mg AUC_0-24_ (A) and AUC_24-48_ (B) versus various continuous infusions (**Supplementary Figures 2-6**).

**FIG 2.**
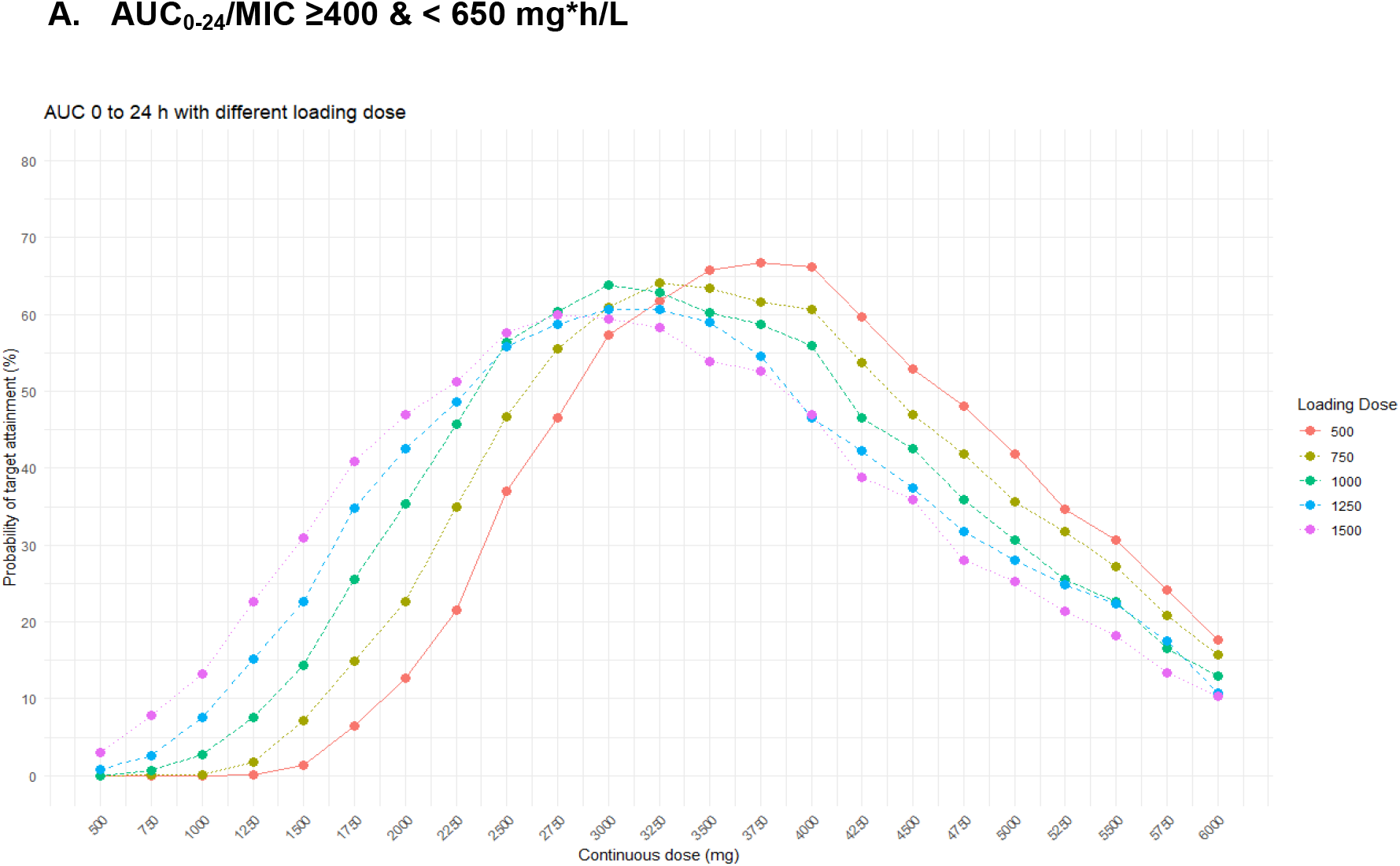
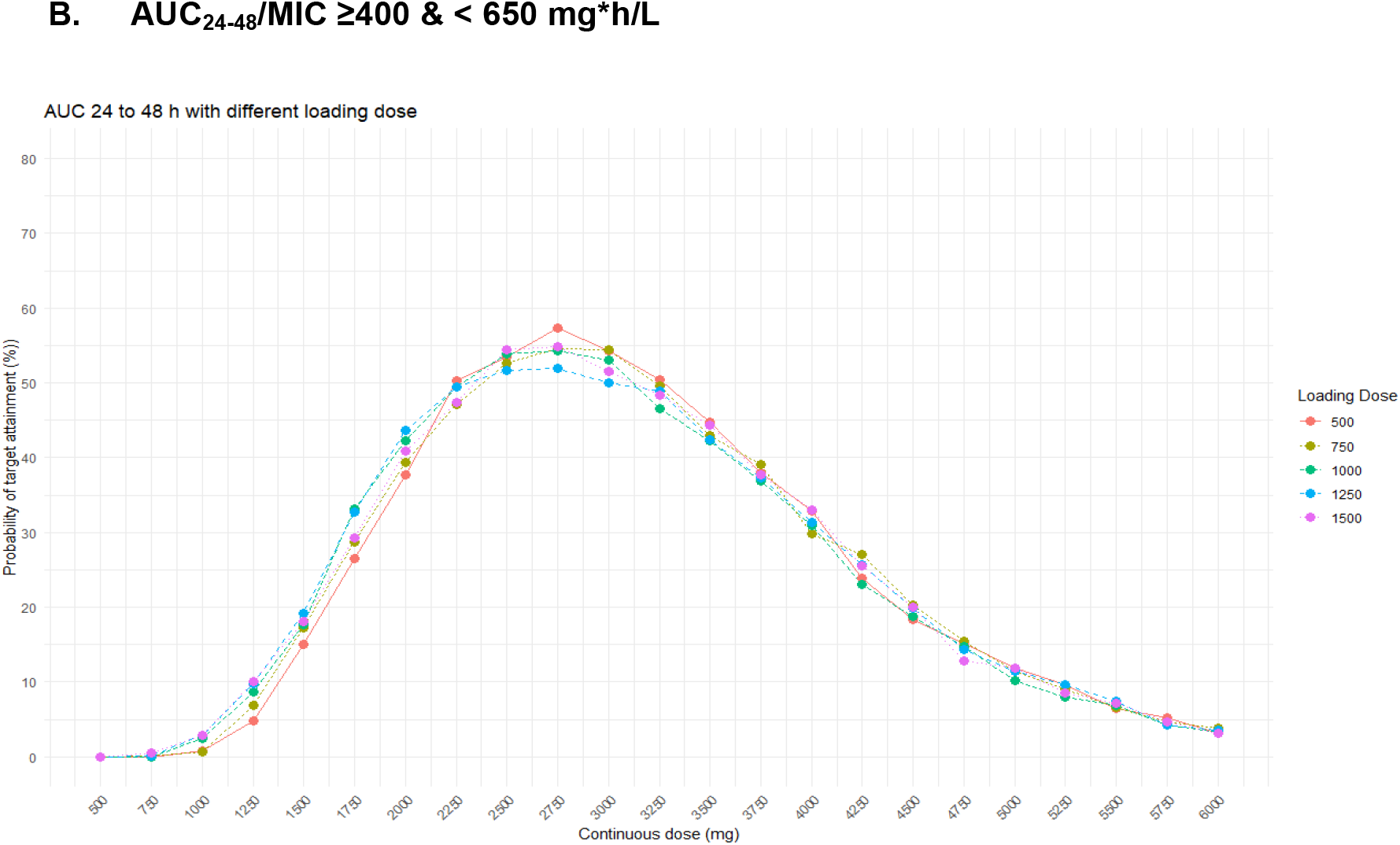
Line and dot plot showing probability of target attainment (%) for efficacy target

## Discussion

The American Thoracic Society Guidelines on Antibiotic Management of Lung Infections recommend IV vancomycin as the first-line treatment of infections due to MRSA. Even though the PK of vancomycin are well studied in other populations (27-29), there is minimal data on the PK of vancomycin in PwCF. Furthermore, the intermittent dosing of vancomycin is not consistent across multiple institutions across the United States, and is dosed at a wide range between 500 and 2,000 mg every 6 -12 hours (6). In this study, we characterized the PK of vancomycin administered as an intermittent and continuous infusion in PwCF with MRSA using a PopPK approach and applied the model to predict the PTA of an efficacy target of AUC/MIC ≥400 mg*h/L and a toxicity target of AUC/MIC <650 mg*h/L. The findings of this study may help inform and optimize initial continuous infusion vancomycin dosing regimens for treating MRSA pulmonary infections in PwCF.

A two-compartment model with proportional residual error and first-order elimination best described the data. The population estimates for CL and Vd were 4.05 L/h and 22.5 L, respectively. Comparing CL and central Vd to peripheral volume, variability was smaller, suggesting a more accurate estimate of these important PK parameters.

The PK of vancomycin in adult PwCF have been previously reported in single-dose studies, as well as in a study of intermittent and continuous infusion dosing. Pleasants et al. evaluated the PK of vancomycin in PwCF after single-dose administration (15 mg/kg), and the PK samples were collected for only up to 24 hours (13). In contrast, we evaluated the PK of vancomycin following intermittent and continuous infusion in PwCF across a period of 2 to 14 days and included a broad range of dosing regimens. In our study, vancomycin levels were at steady state, which is more representative of a clinical situation than the single-dose PK study reported by Pleasants et al. More recently, our group reported the PK of vancomycin in PwCF after intermittent and continuous infusion (14, 15). However, in this current study, we used a more relevant Bayesian estimation approach to calculate the PK parameters. In the pediatric PopPK study by Stockmann et al., the median (range) age of the subjects was 13.9 (8–17) years compared to 27 (19–60) years in our study, indicating that both populations are significantly different from each other (30). We previously reported Po PK of intermittent dosing of vancomycin in adult PwCF and evaluated the PTA of vancomycin for efficacy target of AUC_0– 24_/MIC ≥400 mg*h/L and safety target of AUC_0–24_ <650 mg*h/L after intermittent dosing (16).

The single-dose PK study of vancomycin in PwCF reported a mean Vd of 40.6L and a mean CL of 5.7L/h (13). Our intermittent vancomycin infusion study reported a Vd of 31.5L and a Cl of 5.52L/h. These values are higher than the PK values reported in this study (Vd – 22.5L, CL-4.05/L/h). With intermittent dosing, vancomycin concentrations may vary significantly between the doses. The early rapid distribution overestimates Vd and CL, especially with sparse TDM data. PK estimates following both intermittent and continuous infusion are closer to the real clinical scenario. However, so far, the differences in the effectiveness of continuous and intermittent infusion of vancomycin haven’t been demonstrated in a large-scale study.

Compared to our previous PopPK study of intermittent vancomycin dosing in PwCF, where the highest PTA for an AUC_0-24_/MIC ≥400 and <650 mg·h/L was 65.4% with 2000 mg q12h, our study found higher PTA rates with 3750 mg continuous infusion and 500 mg loading dose (66.7%) (16). Similarly, a PTA of AUC_24-48_/MIC ≥400 mg*h/L & <650 mg*h/L was highest with 2000 mg q12h (64.2%) with intermittent dosing compared to 63.9% with 3000 mg and 57.3% with 2750 mg continuous infusion following the 500 mg loading dose in our study. Thus, continuous infusion can achieve comparable PTA with a lower total vancomycin dose than intermittent infusion. Continuous infusion of vancomycin has been explored as an alternative to intermittent infusion, considering the potential of higher target attainment, less variability in concentration, less need for frequent TDM, and lower risk of nephrotoxicity (19, 31). Fung et al. in 3 case reports demonstrated successful treatment of MRSA in PwCF by switching from intermittent to continuous infusion vancomycin dosing (32).

The estimated PK parameters of vancomycin in our study differed from those in non-critically ill and critically ill populations. Compared to non-critically ill adults, as Nivia et al. (33) reported in a scoping review of 36 PopPK studies, PwCF showed higher CL and lower Vd. On the other hand, CL was almost similar, but a lower Vd was found in our study compared to critically ill adults, where reported CL and Vd were 4.5 (1.3-5.8) L/h and 75.6 (30.1-161) L, respectively (34). The higher CL compared to non-critically ill patients suggests likely increased drug elimination attributed to disease-related factors like altered renal function. These findings highlight the unique PK of vancomycin in PwCF and emphasize the need for personalized dosing strategies to optimize efficacy and safety.

Covariate analysis evaluated the effect of co-administration with inhaled tobramycin and aztreonam on the PK of vancomycin in PwCF. While tobramycin and aztreonam are primarily excreted by the kidneys after inhalation and can influence the PK of vancomycin, coadministration of the two drugs did not significantly impact vancomycin PK. This is expected as systemic absorption of tobramycin and aztreonam was reported to be minimal, and plasma concentrations may not have reached significant levels to alter the PK of vancomycin in PwCF (35, 36). Furthermore, in most hospitals, inhaled antibiotics are not administered during IV antibiotic treatment in PwCF with pulmonary exacerbations (PEx) due to the risk of nephrotoxicity (37). Covariate analysis also demonstrated that CrCL significantly influenced the PK of vancomycin. As vancomycin is eliminated by glomerular filtration, its CL is expected to depend on renal function. Previous studies of continuous vancomycin infusion in critically ill adult patients also found CrCL to be an important predictor of clearance (34). Furthermore, covariate analysis indicated there was not a significant impact of weight on Vd. In addition, the estimated Vd in our analysis was closer to the Vd reported for the non-critically ill population. In general, the impact of weight on the Vd of vancomycin is debatable (34).

Altered physiology in PwCF can significantly affect vancomycin PK. Increased renal clearance, reduced fat mass, chronic inflammation, decreased protein production, and binding can cause large IIV in both CL and Vd (12). A weight-based or empirical dosing approach may prove inadequate, especially for AUC-based targeted regimens. This highlights the need for PopPK analysis specific to PwCF, where loading dose and continuous infusion are needed to maintain stable and predictable drug exposure. A model-informed continuous infusion strategy following a loading dose can help achieve a balance between efficacy and safety goals. Higher target attainment was observed when model-based dosing was used with Bayesian dose adjustment following early TDM (38).

There are some limitations in our study. It was a single-center study, and the PopPK model was developed using retrospective sparse TDM data. The data did not include markers for efficacy, such as MIC of vancomycin against MRSA in the patients included in the study, or forced expiratory volume in 1 second. The efficacy and safety targets (AUC_0-24_/MIC) used for the simulations were calculated using MIC values of 1 mg/L. The PTA can change with the MIC values, which may vary with different centers. Pharmacodynamic (PD) outcome data was not included (clinical or microbiological cure), preventing PK /PD analysis.

Considering the wide range of vancomycin dosing in the clinical setting in the absence of dosing guidelines, future studies should focus on multicenter population models specific to the needs of PwCF receiving intermittent and continuous infusion. Such models can provide a better estimate of vancomycin exposure. In addition, future models can be integrated with Bayesian tools to enable real-time dose adjustment to improve efficacy and safety. PopPK analysis can help bridge the gap in current CF-specific antibiotic dosing guidelines. Future PD studies are needed to establish markers of efficacy and safety, which can be used to guide optimal vancomycin dosing to treat pulmonary exacerbations in PwCF.

## Conclusion

Vancomycin PK after combined intermittent and continuous dosing was adequately described by a two-compartment model with first-order elimination in adult PwCF. The PK parameters Vd and CL in PwCF varied compared to the non-critically ill and critically ill populations, and with intermittent infusion dosing of vancomycin alone. These variations warrant the exploration of intermittent and continuous infusion in large-scale studies in adult PwCF.

## Materials and Methods

### Study population

Data were collected retrospectively from adult PwCF admitted to the University of Utah Hospital between May 2014 and August 2020 for pulmonary exacerbations (PEx). Patients received vancomycin infusion as per the clinician’s discretion, either intermittently, continuously, or both. Patient demographics, including sex, age, weight, height, serum creatinine, and use of concomitant inhaled antibiotics such as tobramycin and aztreonam were collected from the EPIC. Creatinine clearance was calculated using the Cockcroft-Gault formula (20). The body surface area used to normalize creatinine clearance was calculated using the DuBois and DuBois method (21). Patients were excluded if they were pregnant, as determined by serum human chorionic gonadotropin levels. Patients were also excluded if they received IV vancomycin for less than 24 hours. This study received an exemption from the University of Utah Institutional Review Board (IRB #00136598).

### Dosing and sampling schedule

Varying doses of intermittent (range: 500 – 1500 mg) and continuous (range: 1500 - 9000 mg) vancomycin infusion were given to the patients admitted to University of Utah Hospital for an pulmonary exacerbation (PEx). Prior to initiating continuous infusion, patients received a 20 mg/kg loading dose of vancomycin, followed by a continuous 24-hour infusion. Vancomycin continuous doses were administered by IV infusion over a 24-hour duration. The time of random sample collection following continuous infusion varied greatly (7.2-33.1 hours). PwCF were admitted for PEx and received care for 10 to 14 days in the inpatient setting at the University of Utah Hospital. Some PwCF patients had repeat admissions, and data from each admission encounter were included in the data set. Because patients are often co-infected with Pseudomonas aeruginosa, inhaled antibiotics tobramycin and aztreonam were included in the covariate analysis and were administered to patients by nebulization at doses: 300 mg every 12 hours for tobramycin and 75 mg every 8 hours for aztreonam. If the patients were receiving inhaled vancomycin prior to their admission for a PEx, it was held during the hospitalization and therefore not included in the analysis.

### Vancomycin assay

Serum vancomycin concentrations were measured using a Multigent vancomycin assay on Architect cSystem by Abbott Laboratories, IL, USA. The Multigent assay is a homogenous particle-enhanced turbidimetric inhibition immunoassay. The assay was validated for clinical use, and the lower limit of quantitation (LLOQ) for vancomycin assay was 1.1 mg/L. The assay was linear within the vancomycin concentration range of 1.1 to 100 mg/L. The intra- and interday precision for vancomycin assay were 1.38% and 1.52%, respectively. The percentage recovery of vancomycin across a concentration range of 2.5 to 75 mg/L ranged from 99.29% to 105.2%.

### Pharmacokinetic model development

The PopPK analysis for vancomycin plasma concentrations was performed using a non-linear mixed-effect modeling approach using NONMEM® 7.5, ICON plc., MD, USA interfaced with Perl-speaks-NONMEM (PsN^®^) version 5.3.0 (22) and Finch Studio™ NONMEM workbench, Enhanced Pharmacodynamics LLC, NY, USA (23). The pre- and post-processing of the data sets and development of graphics were performed using the Tidyverse package in R software (24). First-order conditional estimation method with interaction was used to estimate the population values of the PK parameters, interindividual variability (IIV), and residual unexplained variability (RUV). Prediction-corrected visual predictive check (pcVPC) was performed with PsN^®^ version 5.3.0.

### Structural and error models

The base structural model was evaluated using different compartment models (e.g., one compartment versus two compartments) with first-order elimination. The predictive performance of the developed PopPK model was determined using diagnostic plots for all observed versus population-predicted vancomycin concentrations and observed versus individual-predicted vancomycin concentrations. Residuals and conditional-weighted residuals were plotted versus time- or population-predicted vancomycin concentrations. Models were further compared by assessing the precision of the parameter estimates, measures of variability, and the objective function value (OFV).

Model variability and random effects were classified as one of two types of errors: interindividual variability and residual unexplained variability. The IIV is the variability inherent between different patients and was assumed to be log-normally distributed according to an exponential equation of the form:

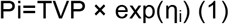

where P_i_ is the individual PK parameter, TVP is the typical population mean of the PK parameter, and η_i_ is the proportional difference between the *i*^th^ subject’s parameters estimate and the typical population mean. Eta is assumed to be normally distributed with a mean of 0 and a variance of ω^2^ (25).

RUV reflects the difference between the model prediction for the individual and the measured observation. This includes errors in the assay, errors in drug dose, errors in the time of measurement, etc. RUV was evaluated using additive, proportional, and combined error models during model development. The proportional error model resulted in the greatest improvement in the OFV. The equation for the proportional error model was

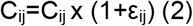

where C_ij_ is the individual observed plasma concentration at time *j*, C_ij_ is the individual-predicted plasma concentration at time *j*, and ε_ij_ is the proportional error, which is assumed to be normally distributed with a mean of 0 and a variance of σ^2^.

The area under the plasma concentration versus time curve over the first 2 days (AUC_0–24_ and AUC_24–48_) for vancomycin was calculated for each patient with linear trapezoidal with linear interpolation method using pracma R package (26).

### Covariate analysis

The covariates evaluated in the analysis were age, weight, creatinine clearance (CrCL), body surface area-normalized creatinine clearance, and concomitant use of inhaled antimicrobials. The inhaled antimicrobials included tobramycin alone, aztreonam alone, or both tobramycin and aztreonam. Race was not included in the covariate analysis, as most of the patients in the data set were white. Sex was not included, as there was no difference in exploratory analysis plots between male and female patients. Further testing was performed by evaluating potential covariates using stepwise forward addition and stepwise backward elimination procedures. A reduction in the OFV of more than 3.84 (*p* < 0.05) was required to retain covariates in the forward addition step. In the backward elimination step, covariates were retained if they reduced the OFV by more than 10.83 (*p* < 0.001). For continuous covariates, age, weight, and CrCL power models were considered:

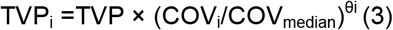

where TVP_i_ represents the individual PK parameter, TVP represents the typical value of the parameters, and θi represents the scale factor for the individual covariate (COV_i_) and the median covariate value COV_median_.

The model for a categorical covariate concomitant inhaled antibiotics was expressed using a proportional model:

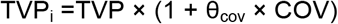

where TVP represents the typical value of the parameters, and θ_cov_ represents the fractional change in the typical parameter with the covariate (COV), where COV has at least two or more levels.

### Model evaluation

The likelihood ratio test (−2LLR) was used to calculate the difference between the nested models, the difference in objective function values (ΔOFV), and between models with a decrease ≥3.84, which is statistically significant at an α = 0.05. Non-nested models were evaluated by using the Akaike information criterion. The final selection of models was also based on goodness-of-fit plots, along with the reliability of the parameter estimates, such as reduced variability between subjects and residual errors. An internal validation method, pcVPC, was performed to evaluate the final model. The original data set and final model parameter estimates were used to simulate 1000 individual datasets. At each time point, the observed median (50th percentile) and the 5th and 95th percentiles were plotted against the corresponding percentiles derived from simulated datasets. The 90% and 95% prediction intervals (PI) around these simulated percentiles were also calculated to assess the model’s performance in capturing central tendency and variability.

### Model simulations

Based on the final PopPK model for vancomycin, Monte Carlo simulations were performed using NONMEM to evaluate the PTA for different vancomycin dosing scenarios (loading dose followed by continuous infusion). The simulated dosing scenarios included a single loading dose given as an intermittent infusion of 500, 750, 1000, 1250, and 1500 mg, followed by various continuous dose regimens ranging from 500 to 6000 mg per 24 hours. The primary PK/PD target for vancomycin included AUC_0-24_/MIC ≥400 - <650 mg*h/L. In total, 23 different continuous dosing scenarios were evaluated. The MIC value of 1 µg/mL was tested for efficacy and safety targets. The PTA was evaluated by determining the percentage of the simulated patient population in which the specified PK/PD target (i.e., AUC_0–24_ and AUC_24–48_ /MIC ≥400 - <650 mg*h/L) was achieved. The PTA was also evaluated by including a toxicity threshold of AUC_0–24_ and AUC_24–48_ ≥650 mg*h/L to the PK/PD target.

## Acknowledgements

This research received no specific grant from any funding agency in the public, commercial, or not-for-profit sectors.

## Conflict of Interest

The authors declare they have no conflict of interest in this work. This work was completed by Rachel Hudson while she was a postdoctoral research fellow at the University of Utah. The Bristol Myers Squibb company was not involved in this study.

